# A Novel Autoantibody against LAG-3 in Systemic Lupus Erythematosus

**DOI:** 10.64898/2026.11.14.761730

**Authors:** Yuya Muramoto, Masahiro Shiokawa, Yukako Maeda, Takeshi Kuwada, Yoshihiro Nishikawa, Tatsuaki Tsuruyama, Takahisa Maruno, Takafumi Yanaidani, Kenji Sawada, Ayako Hirata, Koki Chikugo, Masataka Yokode, Shimpei Matsumoto, Sakiko Ota, Tomoaki Matsumori, Kazushi Izawa, Eitaro Hiejima, Keiji Akamine, Mao Mizuta, Hiroyuki Yoshida, Masaaki Fujita, Ran Nakashima, Hajime Yoshifuji, Tsutomu Chiba, Junko Takita, Akio Morinobu, Hiroshi Seno

## Abstract

Systemic lupus erythematosus (SLE) is an autoimmune disease characterized by immune tolerance loss, although its pathogenesis remains incompletely understood. Immune checkpoint molecules maintain peripheral tolerance, and their dysregulation contributes to autoimmunity. Autoantibodies against lymphocyte activation gene-3 (LAG-3), an immune checkpoint molecule, were identified in a subset of patients with SLE. We enrolled 130 patients with SLE, 119 patients with other autoimmune diseases, and 32 healthy controls. The sera of patients with SLE were screened for antibodies against various immune checkpoint molecules to identify clinically relevant disease-specific autoantibodies against LAG-3. Functional assays were performed to assess the effect of anti-LAG-3 autoantibodies on the interaction between LAG-3 and major histocompatibility complex (MHC) class II. Pathogenic relevance was evaluated using genetic deletion and immunization-based mouse models. Anti-LAG-3 autoantibodies were detected in 52/130 patients with SLE (40.0%) but only 4 of 119 patients with other autoimmune diseases (3.4%). These autoantibodies were associated with lupus nephritis and higher disease activity. In mice, *Lag3* deletion or human LAG-3 immunization induced lupus-like autoimmunity. These findings support the clinical relevance of anti-LAG-3 autoantibodies in SLE and implicate LAG-3 checkpoint disruption in lupus-like autoimmunity.

## INTRODUCTION

Systemic lupus erythematosus (SLE) is a prototypic systemic autoimmune disease characterized by loss of immune tolerance and production of multiple autoantibodies, leading to chronic inflammation and organ damage that commonly affects the skin, joints, kidneys, and central nervous system.^1^ SLE pathogenesis involves complex interactions among genetic, environmental, and immunological factors^1,2^ and remains incompletely understood. Aberrant T- and B-cell activation drives sustained autoantibody production and immune complex deposition in multiple tissues, which are central to SLE pathophysiology.^1,2^

Immune homeostasis depends on the balance between activating and inhibitory signals. Immune checkpoint receptors (ICRs), including cytotoxic T-lymphocyte antigen-4 (CTLA-4), programmed death-1 (PD-1), and lymphocyte activation gene-3 (LAG-3) are inhibitory regulators that restrain lymphocyte activation and maintain self-tolerance.^3–8^ Disruption of these pathways can induce autoimmunity; for example, *Pdcd1*-deficient mice spontaneously develop lupus-like disease.^9^ In humans, therapeutic antibodies targeting ICRs are widely used in cancer immunotherapy, but can trigger immune-related adverse events, affecting multiple organs.^10^

We previously identified an anti-integrin αvβ6 autoantibody as a disease-associated autoantibody in ulcerative colitis using an enzyme-linked immunosorbent assay (ELISA)-based screening strategy.^11^ This finding has since been independently validated in multiple studies.^12–15^ Based on this observation and the immunoregulatory role of ICRs, we hypothesized that patients with SLE may develop autoantibodies against ICRs that impair their inhibitory function and contribute to systemic autoimmunity. To test this hypothesis, we applied the same ELISA-based approach to examine the presence of such autoantibodies in patients with SLE.

## RESULTS

### LAG-3 autoantibodies are present in a subset of patients with SLE

We evaluated the presence of autoantibodies against various immune checkpoint molecules, including PD-1, programmed cell death-ligand 1 (PD-L1), programmed cell death-ligand 2 (PD-L2), CTLA-4, T-cell immunoreceptor with immunoglobulin and ITIM domains (TIGIT), CD96, LAG-3, and T-cell immunoglobulin and mucin domain-containing-3 (TIM-3), in sera of patients with SLE using ELISA. In this preliminary screening, only anti-LAG-3 autoantibodies were detected at a substantial frequency (3/10, 30%), whereas reactivity against other immune checkpoint molecules was negligible (Fig. 1a).

**Fig. 1.**
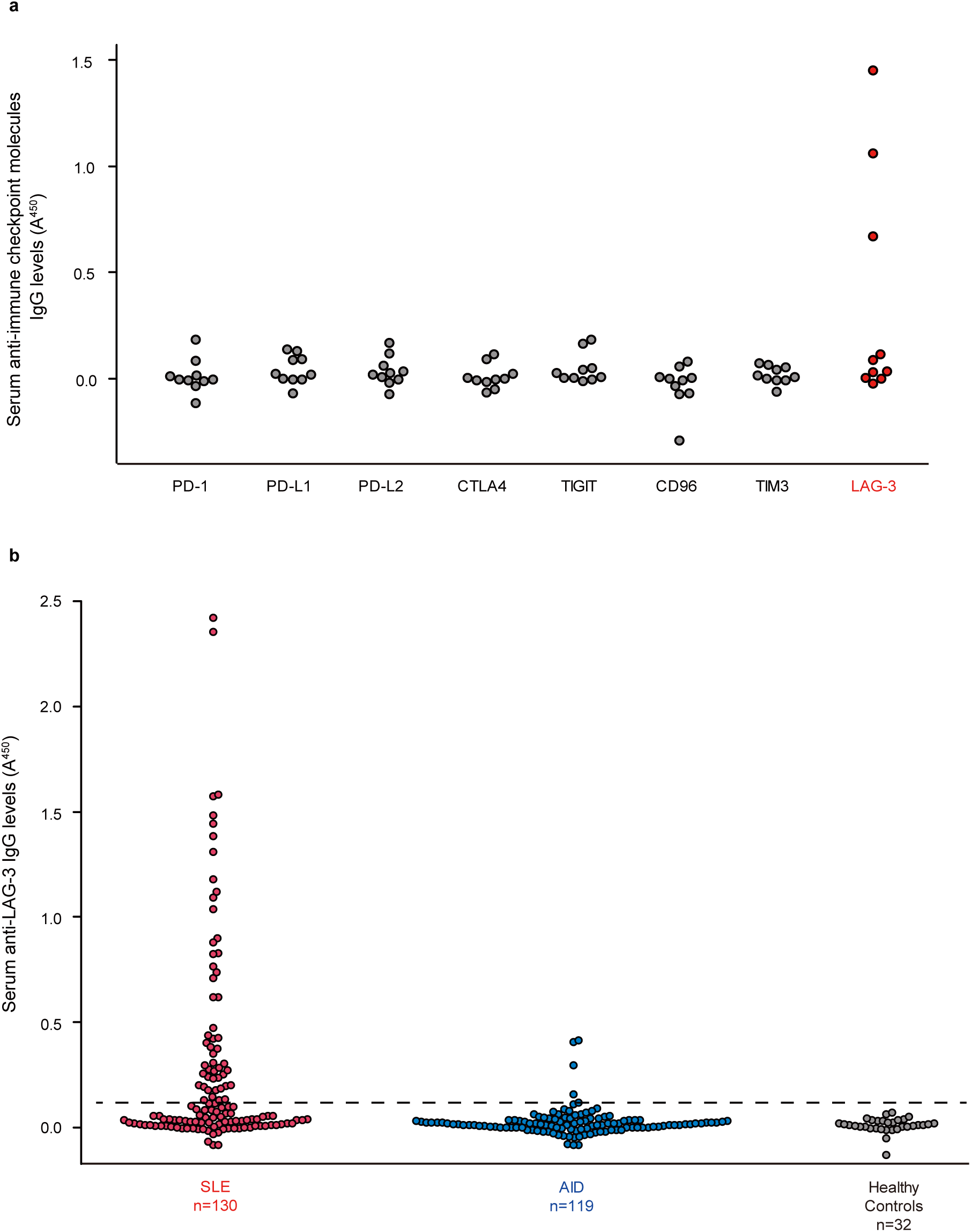
Anti-LAG-3 autoantibodies are present in a subset of patients with SLE. (a) Serum IgG reactivity against multiple immune checkpoint molecules (PD-1, PD-L1, PD-L2, CTLA-4, TIGIT, CD96, TIM-3, and LAG-3) in patients with systemic lupus erythematosus (SLE, n=10) was evaluated using ELISA as a pilot screen. Only anti-LAG-3 autoantibodies were detected at substantial levels. (b) Anti-LAG-3 IgG levels were then measured using ELISA in patients with SLE (n=130), other autoimmune diseases (AIDs, n=119), and healthy controls (n=32). The dashed line indicates the cutoff value, defined as the mean optical density (OD) of healthy controls plus 3 standard deviations (mean + 3SD).

We subsequently expanded the analysis to include 130 patients with SLE, 119 patients with other autoimmune diseases (AIDs), and 32 healthy controls. Using a cutoff value calculated from the 32 healthy controls included in this study (mean+3 standard deviations [SD]; optical density [OD]: 0.119), anti-LAG-3 autoantibodies were detected in 52 of 130 patients with SLE (40.0%), compared with only 4 of 119 patients with AIDs (3.4%) and none of the healthy controls showed positive results (Fig. 1b). Overall, anti-LAG-3 autoantibodies distinguished SLE from other autoimmune diseases with high specificity (96.6%) but moderate sensitivity (40.0%).

### Anti-LAG-3 autoantibodies in patients with SLE mainly belong to the IgG1 subclass

To further characterize anti-LAG-3 autoantibodies, we analyzed IgG subclasses and immunoglobulin isotypes using sera from a representative subset of 40 patients with anti-LAG-3 autoantibody-positive SLE. ELISA revealed that the anti-LAG-3 autoantibodies predominantly belonged to the IgG1 subclass (90%), whereas IgG2, IgG3, and IgG4 subclasses were detected at lower frequencies (5.0%, 5.0%, and 20.0%, respectively; Supplementary Fig. 1a). Other immunoglobulin isotypes, including IgA, IgM, and IgE, were detected only in a small proportion of patients (7.5%, 2.5%, and 2.5%, respectively; Supplementary Fig. 1b).

### Anti-LAG-3 autoantibodies are clinically relevant in patients with SLE

We next investigated the clinical significance of anti-LAG-3 autoantibody positivity by analyzing the patient clinical characteristics (Table 1 and Supplementary Table 1).

**Table 1.** Baseline characteristics of the patients with SLE, according to anti-LAG-3 autoantibody status.

|  |  | **Anti-LAG-3 autoantibody** |  |  |
|---|---|---|---|---|
|  | **All SLE**<br>**n=130** | **Positive**<br>**n=52** | **Negative**<br>**n=78** | **P value<sup>a,b</sup>** |
| **Demographics and Clinical Characteristics** |  |  |  |  |
| Age, median (range) | 46 (18–81) | 42.5 (20–72) | 48 (18–81) | **0.0378** |
| Female sex, n (%) | 115 (88.5) | 48 (92.3) | 67 (85.9) | 0.262 |
| Median time from initial diagnosis of SLE, mo (range) | 108 (0–567) | 115.5 (0–497) | 106 (0–567) | 0.541 |
| SLEDAI-2K score, median (range) | 2 (0–23) | 3 (0–23) | 2 (0–23) | 0.0915 |
| **SLE manifestations, n (%)** |  |  |  |  |
| Fever | 80 (61.5) | 29 (55.8) | 51 (65.4) | 0.270 |
| Mucosal ulcers | 25 (19.2) | 7 (13.5) | 18 (23.1) | 0.173 |
| Malar rash | 73 (56.2) | 27 (51.9) | 46 (59.0) | 0.427 |
| Alopecia | 53 (40.8) | 19 (36.5) | 34 (43.6) | 0.423 |
| Photosensitivity | 61 (46.9) | 20 (38.5) | 41 (52.6) | 0.114 |
| Arthritis | 95 (73.1) | 37 (71.2) | 58 (74.4) | 0.686 |
| Serositis | 26 (20.0) | 10 (19.2) | 16 (20.5) | 0.858 |
| Leukopenia | 86 (66.2) | 31 (59.6) | 55 (70.5) | 0.198 |
| Thrombocytopenia | 44 (33.8) | 8 (15.4) | 36 (46.2) | **0.000281** |
| Nephritis | 61 (46.9) | 31 (59.6) | 30 (38.5) | **0.0180** |
| Neuropsychiatric SLE | 25 (19.2) | 7 (13.5) | 18 (23.1) | 0.173 |
| **Immunological and Serological Profile** |  |  |  |  |
| Anti-double-stranded DNA antibodies, n (%) | 76 (58.5) | 32 (61.5) | 44 (56.4) | 0.561 |
| Anti-Smith antibodies<sup>c</sup>, n (%) | 45 (35.4) | 20 (39.2) | 25 (32.9) | 0.465 |
| Complement C3 <73 mg/dL, n (%) | 46 (35.4) | 18 (34.6) | 28 (35.9) | 0.881 |
| Complement C4 <11 mg/dL, n (%) | 31 (23.8) | 15 (28.8) | 16 (20.5) | 0.275 |
| Total Complement Activity CH50 <30 U/mL, n (%) | 34 (26.2) | 15 (28.8) | 19 (24.4) | 0.568 |
| IgG (mg/dL), median (range) | 1317.5 (435–4554) | 1401 (435–3065) | 1290 (554–4554) | 0.981 |

<sup>a</sup>Comparison between anti-LAG-3 antibody-positive and -negative patients.
<sup>b</sup>P values were calculated using the chi-square test or Mann–Whitney U test as appropriate.
<sup>c</sup>Anti-Sm antibody data were available for 127 patients (51 anti-LAG-3-positive and 76 anti-LAG-3-negative patients).

Anti-LAG-3 autoantibody-positive patients were significantly younger than anti-LAG-3 autoantibody-negative patients (median age: 42.5 vs. 48.0 years, P=0.0378). They were significantly more likely to have lupus nephritis compared with autoantibody-negative patients (59.6% vs. 38.5%, P=0.018; Fig. 2a). Consistently, anti-LAG-3 antibody titers were significantly higher in patients with renal involvement than in those without renal involvement (median OD: 0.13 vs. 0.042, P=0.042; Fig. 2b). Thrombocytopenia was significantly less frequent in the anti-LAG-3 autoantibody-positive group (15.4% vs. 46.2%, P=0.000281).

**Fig. 2.**
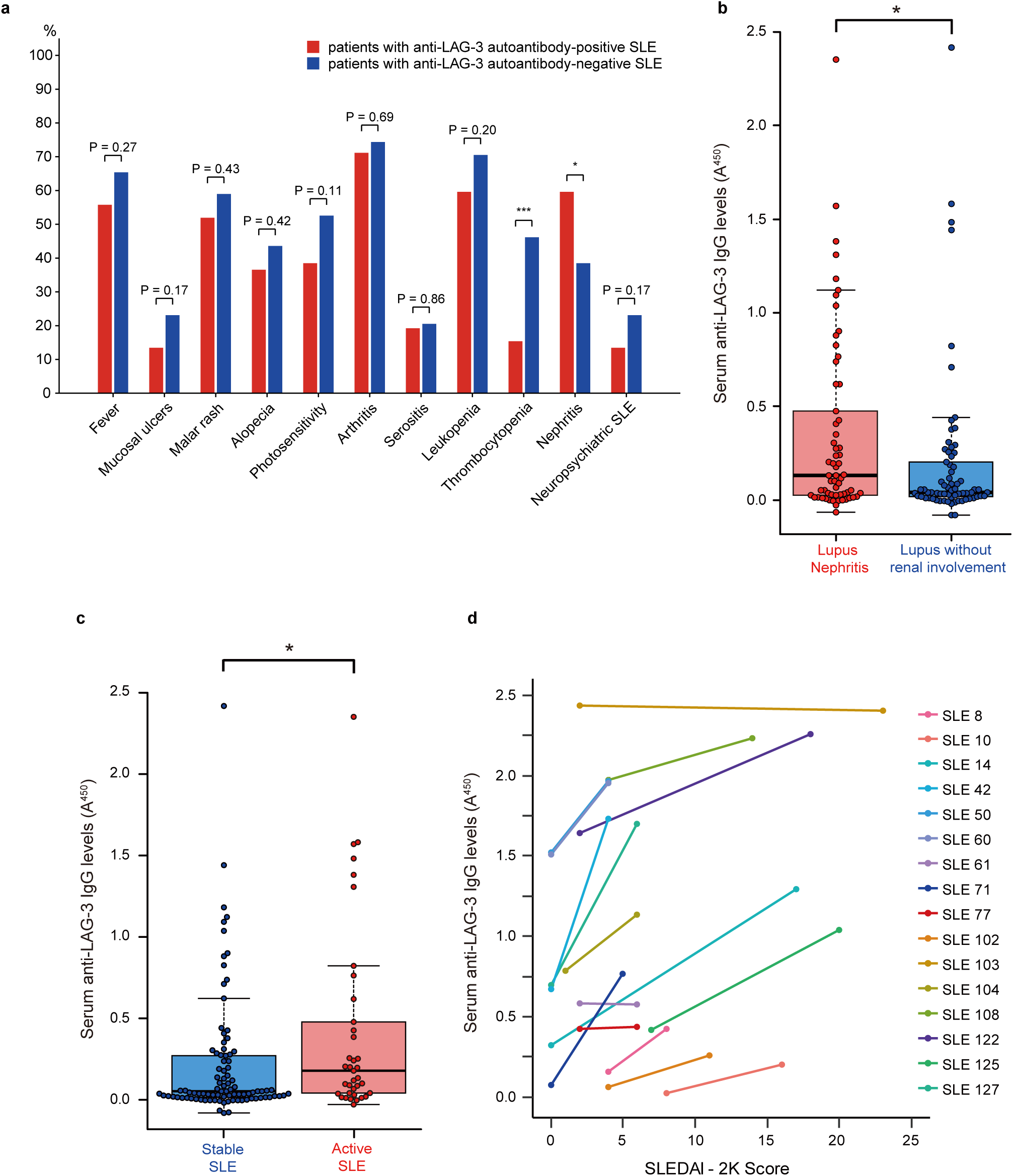
Anti-LAG-3 autoantibodies are clinically relevant in SLE. (a) Frequency of individual clinical manifestations in anti-LAG-3 autoantibody-positive and -negative patients with SLE (n=52 and 78, respectively). (b) Comparison of anti-LAG-3 IgG levels between SLE patients with and without lupus nephritis (n=61 and 69, respectively), as measured using ELISA. (c) Anti-LAG-3 IgG levels in patients with active disease (SLEDAI-2K≥5) and those with stable disease (SLEDAI-2K<5; n=37 and 93, respectively). (d) Paired analysis of serum anti-LAG-3 IgG levels (A450) and SLEDAI-2K scores in selected patients with anti-LAG-3 autoantibody-positive SLE at two time points (n=16). Each line represents an individual patient. Asterisks indicate statistical significance (*P<0.05, ***P<0.001) determined using the chi-square test (panel a) or the Mann–Whitney U test (panels b and c). For box plots, centre lines indicate medians; boxes indicate the interquartile range (IQR); whiskers extend to 1.5× IQR; data points beyond whiskers are shown individually.

Anti-LAG-3 antibody titers were significantly higher in patients with high disease activity (SLE Disease Activity Index 2000 [SLEDAI-2K] ≥ 5) than in those with low disease activity (median OD: 0.177 vs. 0.050, P=0.014; Fig. 2c).

To assess the relationship between anti-LAG-3 IgG levels and disease activity within individuals, we analyzed paired serum samples obtained at two time points with different SLEDAI-2K scores. In 13 of 16 patients, anti-LAG-3 IgG levels were clearly higher at the time point with the higher SLEDAI-2K score, whereas no clear increase was observed in the remaining patients (Fig. 2d). These findings suggest that anti-LAG-3 IgG levels may vary with disease activity within individual patients.

### Anti-LAG-3 autoantibodies inhibit the interaction between LAG-3 and MHC class II

LAG-3 expressed on CD4 T cells maintains peripheral tolerance by binding to MHC class II molecules on antigen-presenting cells and suppressing CD4 T-cell activation.^16^ To determine whether anti-LAG-3 autoantibodies interfere with this immunoregulatory mechanism, we examined purified IgG from six patients each with anti-LAG-3 autoantibody-positive and anti-LAG-3 antibody-negative SLE, and six healthy controls using a luciferase-based LAG-3/MHCII blockade assay (Supplementary Table 3). IgG from anti-LAG-3 autoantibody-positive patients could restore TCR signaling in a dose-dependent manner in four of six cases, whereas IgG from anti-LAG-3 antibody-negative patients and healthy controls showed no effect (Fig. 3).

**Fig. 3.**
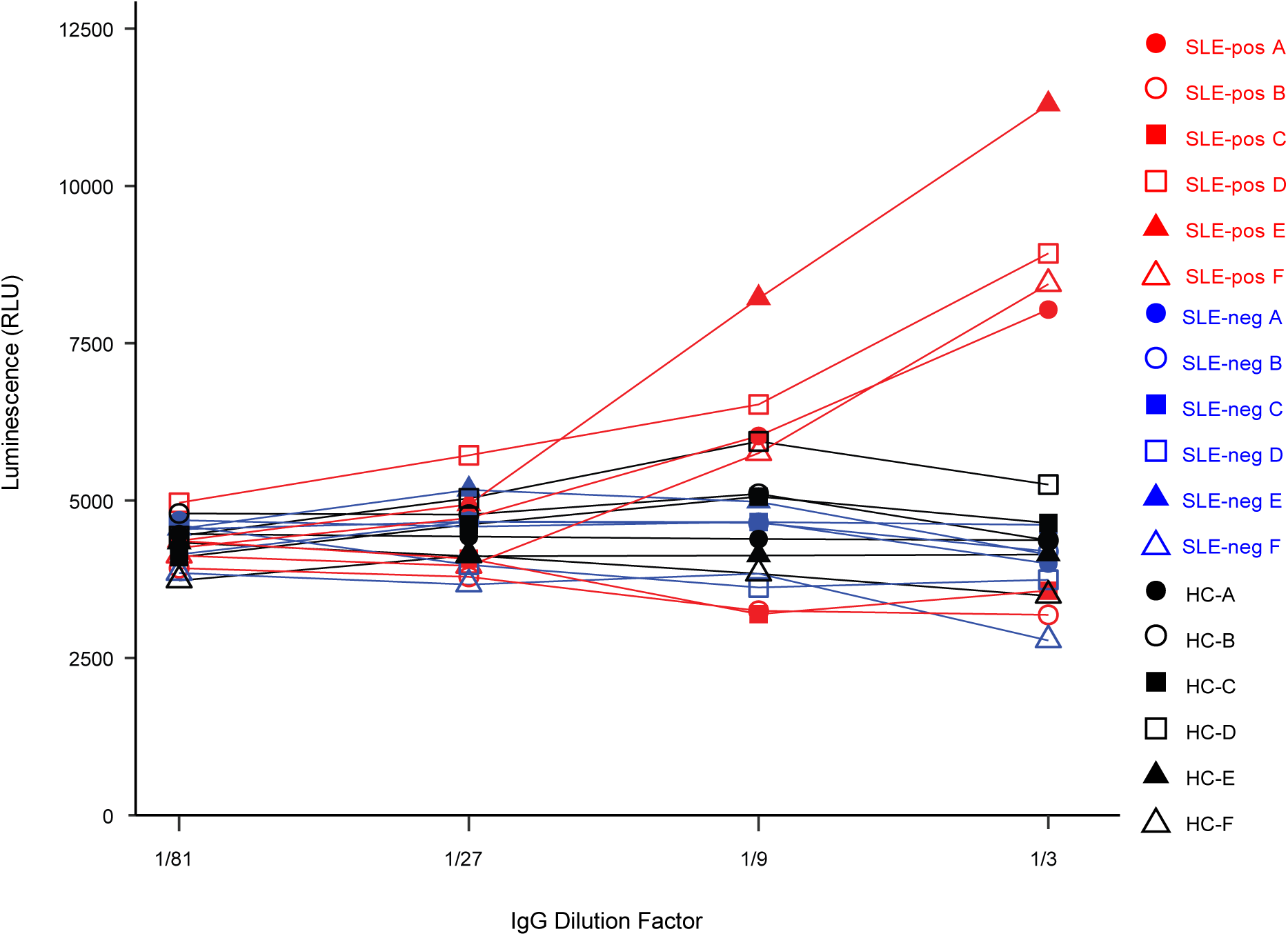
Anti-LAG-3 autoantibodies inhibit the interaction between LAG-3 and MHC class II. Relative light units (RLU) from the LAG-3/MHC class II blockade bioassay are shown for purified IgG from patients with anti-LAG-3 autoantibody-positive SLE (SLE-pos A–F; n=6; red lines), patients with anti-LAG-3 autoantibody-negative SLE (SLE-neg A–F; n=6; blue lines), and healthy controls (HC A–F; n=6; black lines), tested at serial dilutions (1:81, 1:27, 1:9, and 1:3). Increased luminescence reflects disruption of the LAG-3–MHC class II interaction by patient-derived IgG.

These results indicate that anti-LAG-3 autoantibodies disrupt the LAG-3–MHC class II interaction, a critical immune checkpoint pathway; therefore, they may contribute to the breakdown of peripheral tolerance in SLE.

### Lag3-deficient mice produce anti-double-stranded DNA (dsDNA) antibodies as they age and develop glomerulonephritis

As *Pdcd1* knockout mice develop age-dependent glomerulonephritis,^9^ we hypothesized that *Lag3*-deficient mice may exhibit autoimmune features with aging. Therefore, we conducted a 24-month study of global *Lag3* knockout mice. Female *Lag3*-deficient mice developed antinuclear antibodies (ANA) beginning at 12 months of age (Fig. 4a), accompanied by progressively increased anti-dsDNA antibody titers over time (Fig. 4b). Notably, these autoimmune features were predominantly observed in female mice; the data for male mice are provided in Supplementary Fig. 2.

**Fig. 4.**
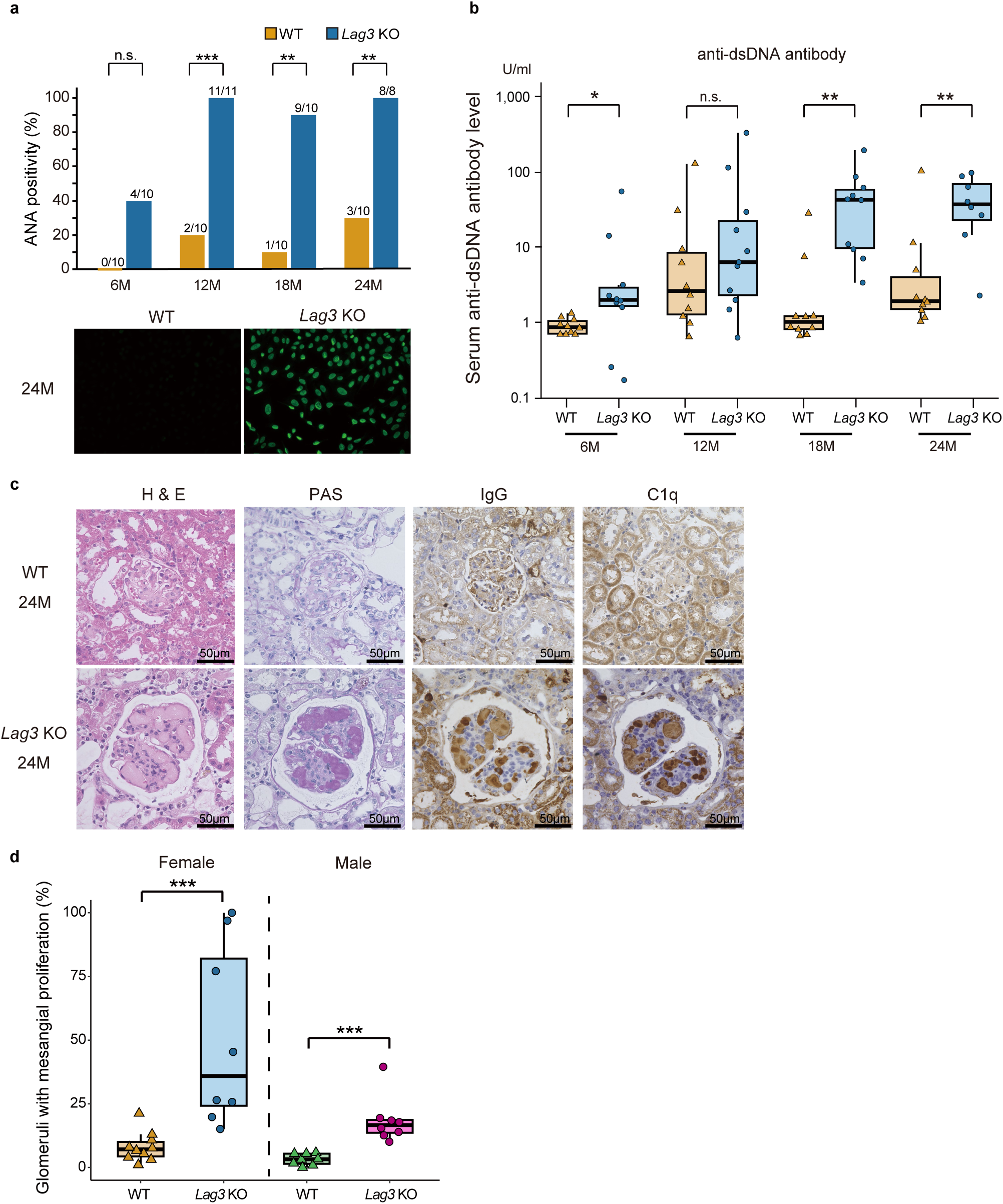
*Lag3*-deficient mice produce anti-dsDNA antibodies as they age and develop glomerulonephritis. (a) Percentage of antinuclear antibody (ANA)-positive female mice at 6, 12, 18, and 24 months of age in wild-type (WT) and *Lag3* knockout (*Lag3* KO) groups. Numbers above the bars indicate the number of ANA-positive mice over the total number of mice assessed at each age. Representative immunofluorescence images of ANA staining at 24 months are shown in the lower panels (WT, left; *Lag3* KO, right). (b) Serum anti-dsDNA IgG levels in female WT and *Lag3* KO mice at each age, measured using ELISA. (c) Representative kidney sections from 24-month-old female WT and *Lag3* KO mice stained with H&E and periodic acid-Schiff stain (PAS), and immunostained for IgG and complement component C1q. Scale bars=50 μm. (d) Quantification of mesangial proliferation in 24-month-old WT and *Lag3* KO mice. The percentage of glomeruli with mesangial proliferation was determined by evaluating all the glomeruli in each kidney section. n.s.=not significant. Asterisks indicate statistical significance (*P<0.05, **P<0.01, ***P<0.001) as determined using Fisher’s exact test (panel a) or the Mann–Whitney U test (panels b and d). For box plots, centre lines indicate medians; boxes indicate the interquartile range (IQR); whiskers extend to 1.5× IQR; data points beyond whiskers are shown individually.

Histological analysis of 24-month-old female *Lag3*-deficient mice revealed features consistent with immune complex-mediated glomerulonephritis, including mesangial expansion and deposition of immune complexes and complement components, resembling lupus nephritis (Fig. 4c). Quantitative evaluation showed that mesangial proliferation was significantly more frequent in *Lag3*-deficient mice than in wild-type controls in both sexes and was more pronounced in females (female: 35.9% vs. 7.1%; male: 16.6% vs. 3.3%, P<0.001 for both; Fig. 4d).

### Mice immunized with human LAG-3 protein show induction of SLE-like lesions

To further investigate whether anti-LAG-3 antibodies contribute to SLE pathophysiology, we immunized female BALB/c mice with recombinant human LAG-3 protein or ovalbumin (OVA) as a control (Fig. 5a).

**Fig. 5.**
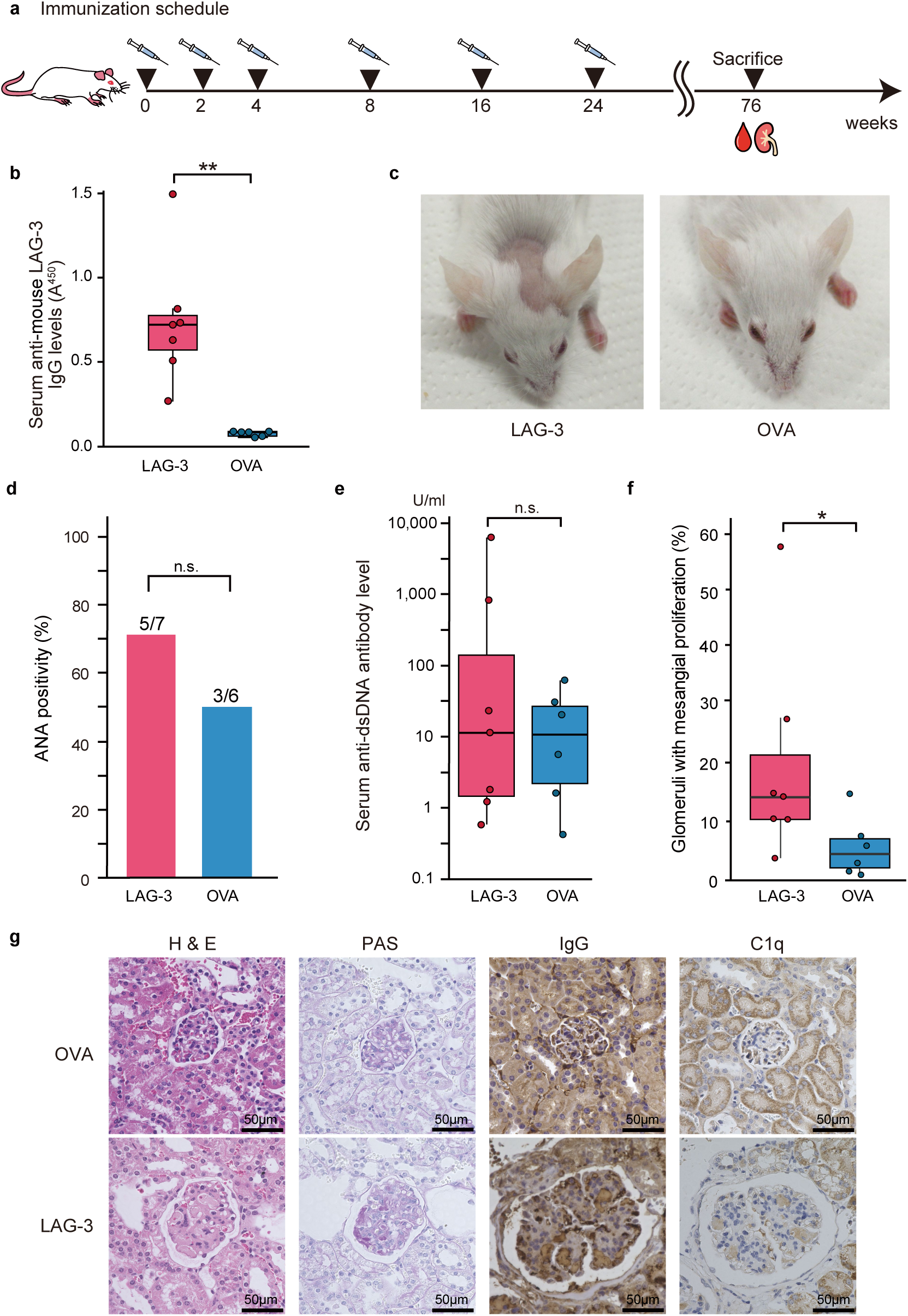
Mice immunized with human LAG-3 protein show induction of SLE-like lesions. (a) Schematic of the immunization protocol: female BALB/c mice were immunized with recombinant human LAG-3 protein or ovalbumin (OVA) as a control, followed by booster injections. (b) Serum anti-mouse LAG-3 IgG antibody levels in LAG-3- and OVA-immunized mice at 12 months post-immunization, measured using ELISA. (c) Representative clinical phenotype showing alopecia on the head of LAG-3-immunized mice, which was absent in OVA controls. (d) Frequency of antinuclear antibody (ANA) positivity in LAG-3- and OVA-immunized mice at 12 months post-immunization. (e) Serum anti-dsDNA IgG antibody levels in LAG-3- and OVA-immunized mice at 12 months post-immunization, measured using ELISA. (f) Quantification of mesangial proliferation in LAG-3- and OVA-immunized mice. The percentage of glomeruli with mesangial proliferation was determined by evaluating all glomeruli in each kidney section. (g) Representative kidney histology in LAG-3- and OVA-immunized mice, showing mesangial proliferation on H&E and PAS staining, and deposition of IgG by immunohistochemistry, which was observed only in LAG-3-immunized mice. Scale bars=50 μm. n.s.=not significant. Asterisks indicate statistical significance (*P<0.05, **P<0.01) as determined using the Mann–Whitney U test (panels b, e and f) or Fisher’s exact test (panel d). For box plots, centre lines indicate medians; boxes indicate the interquartile range (IQR); whiskers extend to 1.5× IQR; data points beyond whiskers are shown individually.

Although the overall sequence identity between human and murine LAG-3 is moderate, the MHC class II-binding interface within the D1 domain is structurally conserved across species.^17^ Considering this conservation and the established practice of using recombinant human proteins to provoke autoimmune responses in mouse models,^18,19^ we chose human LAG-3 as an immunogen to investigate the induction of anti-LAG-3 autoimmunity in mice.

ELISA confirmed that all LAG-3-immunized mice developed anti-mouse LAG-3 antibodies; no such response was observed in the OVA group at 12 months post-immunization (Fig. 5b). Approximately 5 months after immunization, about half of the LAG-3-immunized mice (4/7) exhibited alopecia, predominantly on the top of the head, whereas no clinical changes were observed in the OVA group (0/6; Fig. 5c).

At the time of euthanization, 1 year after the final immunization, the frequency of ANA positivity was higher in the LAG-3 group (5/7) than in the OVA-immunized controls (3/6), though this difference was not statistically significant (Fig. 5d). Similarly, although not statistically significant, some LAG-3-immunized mice showed markedly elevated anti-dsDNA antibody titers compared with OVA-immunized mice (Fig. 5e). Histological examination showed more prominent mesangial proliferation in LAG-3-immunized mice than in the OVA controls. Quantitative analysis confirmed significantly increased mesangial proliferation in the LAG-3-immunized group (14.2% vs 4.4%, P=0.045; Fig. 5f). Moreover, deposition of IgG and the complement component C1q was observed in the kidneys, resembling lupus nephritis (Fig. 5g).

Overall, these observations suggest that disruption of LAG-3 function by LAG-3 autoantibodies contributes to lupus-like disease development in mice.

## DISCUSSION

In this study, we identified anti-LAG-3 autoantibodies selectively in the sera of patients with SLE. They were detected in a substantial proportion of patients with SLE but were largely absent in healthy individuals and patients with other autoimmune diseases, indicating their disease specificity. Functionally, they inhibited the interaction between LAG-3 and MHC class II molecules, a critical checkpoint axis involved in maintaining peripheral tolerance.^16,20–23^ Furthermore, *Lag*3-deficient mice and mice immunized with LAG-3 showed induction of SLE-like lesions. These findings strongly suggest that anti-LAG-3 autoantibodies in patients with SLE inhibit the interaction between LAG-3 and MHC class II molecules and contribute to the breakdown of peripheral tolerance, potentially inducing lupus-like autoimmunity.

These autoantibodies predominantly belonged to the IgG1 subclass, similar to that reported for major SLE autoantibodies, including anti-dsDNA and anti-Smith (Sm) antibodies.^24^ Clinically, anti-LAG-3 antibody positivity was associated with lupus nephritis and higher SLEDAI scores, supporting its relevance to active disease. Although numerous autoantibodies have been described in patients with SLE, their diagnostic performance varies substantially. Anti-dsDNA and anti-Sm antibodies are highly specific but lack sensitivity, whereas anti-Sjögren’s syndrome type A (SSA) or anti-ribonucleoprotein (RNP) are more sensitive but not disease-specific. Thus, anti-LAG-3 antibodies were highly specific for SLE, albeit modestly sensitive, suggesting that they help distinguish SLE from other autoimmune diseases. Their limited sensitivity may reflect the clinical and immunological heterogeneity of SLE. Furthermore, our cohort included several patients with long disease duration (median, 108 months) and relatively low disease activity. These factors may have reduced the detection rate of anti-LAG-3 antibodies for SLE. Anti-LAG-3 antibody positivity was associated with a higher prevalence of nephritis but a lower frequency of thrombocytopenia. Although nephritis and thrombocytopenia often coexist in SLE,^25^ thrombocytopenia may be less frequent among patients with lupus nephritis.^26^ Our findings may be compatible with this observation. Although the reason for this discrepancy remains unclear, a possible explanation is that anti-LAG-3 antibodies preferentially contribute to immune pathways driving renal involvement, whereas thrombocytopenia is often mediated by LAG-3-independent mechanisms.

Using a cell-based blockade assay in this study, we demonstrated that anti-LAG-3 antibodies interfered with the interaction between LAG-3 and MHC class II molecules. A previous study has shown that LAG-3 preferentially binds stable peptide–MHC class II complexes presenting self-antigens and suppresses CD4 T-cell activation through conformation-dependent recognition of these complexes.^16^ In this context, blockade of the LAG-3–MHC class II interaction by anti-LAG-3 antibodies may impair a critical checkpoint that normally restrains CD4 T-cell responses to self-antigens presented by MHC class II. Because aberrant MHC class II-mediated presentation of nuclear self-antigens is a central feature of SLE, disruption of this pathway could promote activation of autoreactive CD4 T cells. Although antigen specificity was not directly examined *in vivo*, interference with the LAG-3–MHC class II axis by anti-LAG-3 antibodies could contribute to the loss of peripheral tolerance in SLE.

*Lag3* deficiency exacerbates autoimmunity when combined with other checkpoint disruptions, manifested in severe myocarditis in *Lag3* and *Pdcd1* double-knockout mice and accelerated onset of type 1 diabetes in *Lag3*-deficient NOD mice.^8,27^ These observations suggest that loss of LAG-3 function can predispose to autoimmunity, although the consequences of *Lag3* deficiency alone over time remain unclear. *In vivo*, we examined two murine models. Aged *Lag3*-deficient mice spontaneously developed lupus-like features, including presence of anti-dsDNA antibodies and glomerular immune complex deposition. These phenotypes were more pronounced in female mice, consistent with the female predominance of SLE. Although disease onset was delayed, these results support a model wherein sustained loss of LAG-3 function promotes immune dysregulation. In parallel, immunization of BALB/c mice with recombinant human LAG-3 induced anti-LAG-3 autoantibodies and led to ANA positivity and glomerular immune complex deposition, even in the absence of known genetic susceptibility. Together, these experimental results indicate that anti-LAG-3 antibodies are not just a consequence of immune activation but may actively contribute to the breakdown of tolerance and development of lupus-like autoimmunity.

In autoimmune diseases, mutations in target autoantigens often mimic the phenotypes induced by pathogenic autoantibodies. For example, loss-of-function mutations in acetylcholine receptor (AChR) subunits cause myasthenic symptoms resembling those induced by AChR autoantibodies in patients with myasthenia gravis.^28,29^ Likewise, a rare germline mutation in *Lag3* has been reported in familial SLE,^30^ indicating that interference with LAG-3 function, whether genetic or antibody-mediated, may contribute to SLE pathophysiology.

This study had several limitations. First, although we detected anti-LAG-3 autoantibodies in patients with SLE and demonstrated their ability to inhibit LAG-3–MHC class II interaction *in vitro*, we did not directly assess their functional effects on T-cell activity in patient-derived immune cells. Further studies using *ex vivo* T-cell assays or single-cell transcriptomic profiling may elucidate the immunological impact of these antibodies.

Second, although our mouse models recapitulated several pathological features of SLE, the immunization experiments employed recombinant human LAG-3 protein, which may not fully reflect the immunogenic properties or epitope presentation of endogenous murine LAG-3. Accordingly, the translational relevance of these models should be interpreted cautiously. In addition, although lupus-like phenotypes in *Lag3*-deficient mice were more pronounced in females, the immunization experiments were performed exclusively in female mice. Therefore, potential sex-specific effects of LAG-3 immunization were not evaluated in this study.

Third, this study was conducted exclusively in a Japanese cohort. Considering the well-established genetic and immunological heterogeneity of SLE across populations, further validation in multiethnic cohorts is necessary to determine the generalizability of anti-LAG-3 autoantibody prevalence and its clinical associations.

Finally, although paired analyses were performed in a small subset of patients, we did not conduct systematic longitudinal sampling across the cohort or prospectively evaluate changes in the anti-LAG-3 autoantibody levels in relation to treatment. Therefore, prospective studies with serial sampling are needed to define their utility as dynamic biomarkers.

In conclusion, our findings identify that anti-LAG-3 autoantibodies are specific to patients with SLE and are functionally active in its pathophysiology, as they interfere with a key immune checkpoint pathway. These results suggest that immune checkpoint dysregulation may contribute to the loss of self-tolerance in SLE. Future studies should investigate the longitudinal dynamics of anti-LAG-3 antibody production, their potential utility as biomarkers of disease activity, and their suitability as targets for therapeutic intervention.

## ONLINE METHODS

### Human subjects and sample collection

We enrolled 130 patients with SLE, 119 patients with other AIDs (Supplementary Tables 1 and 2), and 32 healthy controls. SLE was diagnosed according to one or more of the following guidelines: the 1997 revised American College of Rheumatology (ACR),^31,32^ the Systemic Lupus Erythematosus International Collaborating Clinics (SLICC),^33^ or the most recent ACR/European League Against Rheumatism (EULAR) criteria.^34^ Other AIDs were diagnosed using the standard classification criteria appropriate for each disease.

For immunoglobulin subclass and isotype analyses, 40 patients with anti-LAG-3 autoantibody-positive SLE and 32 healthy controls were selected. Paired serum samples with corresponding SLEDAI-2K scores were available for 16 patients with anti-LAG-3-positive SLE and were used for within-patient analyses.

Serum samples were aliquoted and stored at −80 °C until analysis. Clinical information, including serological markers, laboratory test results, and disease manifestations, was retrospectively collected from medical records (Supplementary Tables 1 and 2).

All participants provided written informed consent, and the study was approved by the Ethics Committee of Kyoto University Graduate School and Faculty of Medicine (protocol number R1004 and R2831).

### Lag3 knockout mice

*Lag3*-deficient mice (B6.129S2-*Lag3^tm1Doi^*/J; The Jackson Laboratory, Stock No: 026644)^35^ were maintained under specific pathogen-free conditions. Female and male mice aged up to 24 months were used. Serum samples were collected at the time of euthanization for detecting ANA by indirect immunofluorescence using HEp-2 cells, and anti-dsDNA antibodies using ELISA.

Kidneys were harvested, fixed in 10% neutral phosphate-buffered formalin, embedded in paraffin, and sectioned at 3-μm thickness. The sections were stained with hematoxylin and eosin (H&E) and periodic acid-Schiff (PAS) reagents for histological analysis.

For immunohistochemistry, paraffin-embedded kidney sections were deparaffinized, rehydrated, and subjected to antigen retrieval by microwaving in either 10 mM citrate buffer (pH 6.0) or EDTA buffer (pH 9.0) for 15 min. Endogenous peroxidase activity was quenched using 3% hydrogen peroxide. After blocking with a commercial blocking solution (X0909; Dako), the sections were incubated overnight at 4 °C with the primary antibodies. The following primary antibodies were used: anti-mouse IgG (1:200 dilution; ab6789, Abcam) and anti-C1q antibody [7H8] (1:200 dilution; ab11861, Abcam). Antibody binding was detected using the VECTASTAIN Elite ABC-HRP Kit (Vector Laboratories), and color was developed using the Liquid DAB+ Substrate Chromogen System (K3468; Dako). All sections were counterstained with hematoxylin and mounted for microscopic evaluation.

### Immunization model

Eight-week-old female BALB/c mice (Japan SLC, Shizuoka, Japan) were immunized subcutaneously with 10 μg of recombinant human LAG-3 protein (LA3-H5222; ACROBiosystems) or ovalbumin (015-24731; FUJIFILM Wako Pure Chemical Corporation), which was emulsified at 1:1 in Complete Freund’s Adjuvant (263810; BD Biosciences). Booster immunizations were administered on days 14 and 28, followed by three additional boosters: one at 1 month and two more at 2-month intervals, using Incomplete Freund’s Adjuvant (011-09551, FUJIFILM Wako Pure Chemical Corporation). Serum and kidney tissues were collected 1 year after the final immunization.

### ELISA for autoantibody detection

Candidate autoantigens were evaluated using human recombinant immune checkpoint proteins obtained from ACROBiosystems (TM3-H5229, LAG-3 LA3-H5222, TAE-H52H0, TIT-H52H3, PD1-H5221, PD1-H5229, PD2-H82E8, CT4-H5229). Serum IgG antibodies against these proteins were measured using the ELISA Starter Accessory Kit (E101; Bethyl Laboratories) following the manufacturer’s protocol and previously published methods.^11,36^

Briefly, microtiter plates were coated with 100 μL of 2 μg/mL recombinant proteins, incubated overnight at 4 °C, blocked, and incubated with 100 µL of diluted serum (1:100) from patients for 60 min at 25 °C. The contents of the plates were washed and then incubated with 100 µL rabbit anti-human IgG antibody conjugated with horseradish peroxidase (HRP) (1:50,000; ab6759, Abcam) at 20–25 °C for 60 min. After rewashing, the bound reactants were detected by incubation with 3,3′,5,5′-tetramethylbenzidine (TMB) for 7 min. Absorbance was measured at 450 nm. To account for nonspecific binding, parallel wells without antigen coating were included for each serum sample, and background absorbance values obtained from these uncoated wells were subtracted from the corresponding antigen-coated well readings to derive the final values.

### IgG subclass and immunoglobulin isotype analysis

To examine the IgG autoantibody subclasses, we used the following secondary antibodies: anti-human IgG1, IgG2, IgG3, and IgG4 conjugated with HRP (1:2,000; A10648, 1:1,000; 05-0520, 1:1,000; 05-3620, and 1:2,000; A106454, respectively; Invitrogen). To examine the isotypes of the autoantibodies, the following secondary antibodies were used: anti-human IgA, IgM, and IgE conjugated with HRP (1:50,000 A80-102P, 1:100,000 A80-100P, and 1:1,000 A80-108P, respectively; Bethyl Laboratories).

### LAG-3/MHCII blockade assay

The functional capacity of anti-LAG-3 autoantibodies to disrupt the interaction between LAG-3 and MHC class II was evaluated using the LAG-3/MHCII Blockade Bioassay (JA1111; Promega) according to the manufacturer’s instructions. This reporter assay uses the principle that LAG-3 engagement with MHC class II molecules on antigen-presenting cells inhibits T cell receptor (TCR) signaling, resulting in decreased luciferase activity. Interference with this interaction restores TCR signaling, leading to increased luminescence.

For this assay, serum samples were obtained from six patients with anti-LAG-3 autoantibody-positive SLE (SLE-pos A–F), six patients with anti-LAG-3 autoantibody-negative SLE (SLE-neg A–F), and six healthy controls (HC A–F)(Supplementary Table 3). The anti-LAG-3 autoantibody-positive patients were selected based on high antibody titers at the disease onset. Most of these SLE samples were obtained from patients outside the main cross-sectional cohort that was analyzed for prevalence and clinical features.

IgG was purified from each serum sample using a protein A affinity resin (P-014; ProteNova). Eluates were buffer-exchanged with PBS and concentrated. IgG concentrations were not normalized between samples, and each sample was tested at its respective native concentration.

### Quantitative assessment of mesangial proliferation

Kidney tissues were fixed in 10% neutral-buffered formalin, embedded in paraffin, sectioned at 3 µm, and stained with PAS reagents. All glomeruli in each section were examined. Mesangial proliferation was defined as the presence of more than three nuclei within a single mesangial area delineated by the mesangial matrix. For each mouse, the percentage of glomeruli meeting this criterion was calculated as the mesangial proliferation index.

### Detection of antinuclear antibodies (ANA)

HEp-2 cells (5 × 10 cells/well; RCB1889, RIKEN BRC CELL BANK) were seeded onto chamber slides (SCS-N08; Matsunami Glass) one day before staining and incubated overnight to allow adherence.

Cells were then fixed in cold acetone (−20 °C, 10 min), washed with phosphate-buffered saline (PBS), and blocked with protein blocking solution (X0909; Dako) for 30 min at 20–25 °C.

Mouse serum was diluted 1:80 in PBS and applied as the primary antibody, followed by overnight incubation at 4 °C in a humidified chamber. After washing, Alexa Fluor 488-conjugated goat anti-mouse IgG (1:400; A32723, Thermo Fisher Scientific) was used as the secondary antibody and incubated for 1 h in the dark at 20–25 °C.

Slides were mounted with a glycerol-based, hard-setting mountant (P36982; Thermo Fisher Scientific), overlaid with 24 × 50 mm coverslips, and cured at 20–25 °C for 24 h before fluorescence microscopy.

### ELISA for detection of anti-mouse LAG-3 antibodies in immunized mice

To confirm the induction of anti-mouse LAG-3 antibodies in immunized mice, serum IgG reactivity against mouse LAG-3 was assessed by ELISA. Recombinant mouse LAG-3/CD223 protein (LA3-M52H5; ACROBiosystems) was used as the coating antigen. Microtiter plates were coated with 100 μL of recombinant mouse LAG-3 protein (2 μg/mL) and incubated overnight at 4 °C. After blocking, plates were incubated with mouse serum diluted 1:100 for 60 min at 20–25 °C. After washing, bound IgG was detected using goat anti-mouse IgG conjugated with HRP (1:50,000; ab6789, Abcam) for 60 min at 20–25 °C. After washing, signal was developed using TMB substrate for 20 min, and absorbance was measured at 450 nm.

### Measurement of anti-ds DNA antibodies in mouse serum

Serum levels of anti**-**dsDNA IgG antibodies were measured using a commercially available ELISA kit (297-91801; also available as AKRDD-061; FUJIFILM Wako) according to the manufacturer’s instructions. Serum samples were initially diluted 1:100 with the supplied sample diluent. For samples with absorbance values outside the linear range of the standard curve, serial dilutions (1:50 for low absorbance, and 1:1,000, or 1:10,000 for high absorbance) were performed as necessary to ensure quantification within the dynamic range of the assay. The concentration of anti-dsDNA antibodies was calculated in units per milliliter (U/mL) based on the standard curve generated in each plate.

### Statistical analysis

All statistical analyses were performed using R (version 4.1.2). Continuous variables were compared using the Mann–Whitney U test. Categorical variables were analyzed using the chi-square test or Fisher’s exact test, as appropriate. P values <0.05 were considered statistically significant. Data are presented as median with range as appropriate.

## DATA AVAILABILITY STATEMENT

The data supporting the findings of this study are available from the corresponding author upon reasonable request, subject to institutional and ethical regulations. Individual-level human data cannot be made publicly available due to privacy and informed consent restrictions.

## ACKNOWLEDGMENTS

We thank the patients and healthy volunteers who provided the serum samples for this study. We are also grateful to Shino Yamaguchi and Taichi Ito for providing excellent technical assistance.

This work was supported by the Agency for Medical Research and Development (AMED; 25ym0126808j0004). The authors also thank Editage (www.editage.com) for English language editing.

## AUTHOR CONTRIBUTION

Yuya Muramoto and Masahiro Shiokawa conceptualized the study. Yuya Muramoto, Yukako Maeda, Tatsuaki Tsuruyama, and Takahisa Maruno performed the investigation. Yukako Maeda, Kazushi Izawa, Eitaro Hiejima, Keiji Akamine, Mao Mizuta, Hiroyuki Yoshida, Masaaki Fujita, Ran Nakashima, and Hajime Yoshifuji contributed resources. Yuya Muramoto wrote the original draft of the manuscript. Yuya Muramoto, Masahiro Shiokawa, Takeshi Kuwada, Yoshihiro Nishikawa, Takafumi Yanaidani, Kenji Sawada, Ayako Hirata, Koki Chikugo, Masataka Yokode, Shimpei Matsumoto, Sakiko Ota, Tomoaki Matsumori, Junko Takita, and Akio Morinobu reviewed and edited the manuscript. Masahiro Shiokawa acquired funding. Masahiro Shiokawa, Tsutomu Chiba, and Hiroshi Seno supervised the study. All authors reviewed the manuscript and approved the final version.

## REFERENCES

1 Tsokos, G. C. Systemic lupus erythematosus. N Engl J Med 365, 2110–2121 (2011). 10.1056/NEJMra1100359.

2 Zhang, Q. & Vignali, D. A. A. Co-stimulatory and co-inhibitory pathways in autoimmunity. Immunity 44, 1034–1051 (2016). 10.1016/j.immuni.2016.04.017.

3 Linsley, P. S. et al. CTLA-4 is a second receptor for the B cell activation antigen B7. J Exp Med 174, 561–569 (1991). 10.1084/jem.174.3.561.

4 Linsley, P. S. et al. Immunosuppression in vivo by a soluble form of the CTLA-4 T cell activation molecule. Science 257, 792–795 (1992). 10.1126/science.1496399.

5 Okazaki, T., Chikuma, S., Iwai, Y., Fagarasan, S. & Honjo, T. A rheostat for immune responses: the unique properties of PD-1 and their advantages for clinical application. Nat Immunol 14, 1212–1218 (2013). 10.1038/ni.2762.

6 Workman, C. J. & Vignali, D. A. A. The CD4-related molecule, LAG-3 (CD223), regulates the expansion of activated T cells. Eur J Immunol 33, 970–979 (2003). 10.1002/eji.200323382.

7 Workman, C. J. & Vignali, D. A. A. Negative regulation of T cell homeostasis by lymphocyte activation gene-3 (CD223). J Immunol 174, 688–695 (2005). 10.4049/jimmunol.174.2.688.

8 Okazaki, T. et al. PD-1 and LAG-3 inhibitory co-receptors act synergistically to prevent autoimmunity in mice. J Exp Med 208, 395–407 (2011). 10.1084/jem.20100466.

9 Nishimura, H., Nose, M., Hiai, H., Minato, N. & Honjo, T. Development of lupus-like autoimmune diseases by disruption of the PD-1 gene encoding an ITIM motif-carrying immunoreceptor. Immunity 11, 141–151 (1999). 10.1016/s1074-7613(00)80089-8.

10 Suijkerbuijk, K. P. M., van Eijs, M. J. M., van Wijk, F. & Eggermont, A. M. M. Clinical and translational attributes of immune-related adverse events. Nat Cancer 5, 557–571 (2024). 10.1038/s43018-024-00730-3.

11 Kuwada, T. et al. Identification of an anti-integrin αvβ6 autoantibody in patients with ulcerative colitis. Gastroenterology 160, 2383–2394.e2321 (2021). 10.1053/j.gastro.2021.02.019.

12 Livanos, A. E. et al. Anti-integrin αvβ6 autoantibodies are a novel biomarker that antedate ulcerative colitis. Gastroenterology 164, 619–629 (2023). 10.1053/j.gastro.2022.12.042.

13 Yang, J., Huang, M. M. C., Liang, M. M. J. & Lei, M. M. Y. The diagnostic performance of serum αvβ6 autoantibodies for ulcerative colitis: a systematic review and meta-analysis. Clin Res Hepatol Gastroenterol 48, 102317 (2024). 10.1016/j.clinre.2024.102317.

14 Muramoto, Y. et al. Anti-integrin αvβ6 antibody as a diagnostic marker for pediatric patients with ulcerative colitis. Gastroenterology 163, 1094–1097.e1014 (2022). 10.1053/j.gastro.2022.06.026.

15 Okabe, M. et al. Anti-integrin αvβ6 antibody as a biomarker for diagnosing ulcerative colitis: a nationwide multicenter validation study. J Gastroenterol 60, 86–95 (2025). 10.1007/s00535-024-02176-x.

16 Maruhashi, T. et al. LAG-3 inhibits the activation of CD4+ T cells that recognize stable pMHCII through its conformation-dependent recognition of pMHCII. Nat Immunol 19, 1415–1426 (2018). 10.1038/s41590-018-0217-9.

17 Ming, Q. et al. Structural basis for mouse LAG3 interactions with the MHC class II molecule I-Ab. Nat Commun 15, 7513 (2024). 10.1038/s41467-024-51930-5.

18 Tseng, C. E. et al. The 52-kd protein as a target of intermolecular spreading of the immune response to components of the SS-A/Ro-SS-B/La complex. Arthritis Rheum 40, 936–944 (1997). 10.1002/art.1780400523.

19 Deshmukh, U. S. et al. Immune responses to Ro60 and its peptides in mice. I. The nature of the immunogen and endogenous autoantigen determine the specificities of the induced autoantibodies. J Exp Med 189, 531–540 (1999). 10.1084/jem.189.3.531.

20 Aggarwal, V., Workman, C. J. & Vignali, D. A. A. LAG-3 as the third checkpoint inhibitor. Nat Immunol 24, 1415–1422 (2023). 10.1038/s41590-023-01569-z.

21 Blackburn, S. D. et al. Coregulation of CD8+ T cell exhaustion by multiple inhibitory receptors during chronic viral infection. Nat Immunol 10, 29–37 (2009). 10.1038/ni.1679.

22 Woo, S. R. et al. Immune inhibitory molecules LAG-3 and PD-1 synergistically regulate T-cell function to promote tumoral immune escape. Cancer Res 72, 917–927 (2012). 10.1158/0008-5472.can-11-1620.

23 Mariuzza, R. A., Shahid, S. & Karade, S. S. The immune checkpoint receptor LAG3: Structure, function, and target for cancer immunotherapy. J Biol Chem 300, 107241 (2024). 10.1016/j.jbc.2024.107241.

24 Eisenberg, R. A., Dyer, K., Craven, S. Y., Fuller, C. R. & Yount, W. J. Subclass restriction and polyclonality of the systemic lupus erythematosus marker antibody anti-Sm. J Clin Invest 75, 1270–1277 (1985). 10.1172/jci111826.

25 Ktona, E. et al. Evaluation of thrombocytopenia in systemic lupus erythematosus and correlation with different organs damages. Mater Sociomed 26, 122–124 (2014). 10.5455/msm.2014.26.122-124.

26 Diao, H. et al. Clinicopathological characteristics and outcomes of lupus nephritis patients with thrombocytopenia: a single-center retrospective study. Immun Inflamm Dis 13, e70179 (2025). 10.1002/iid3.70179.

27 Bettini, M. et al. Cutting edge: accelerated autoimmune diabetes in the absence of LAG-3. J Immunol 187, 3493–3498 (2011). 10.4049/jimmunol.1100714.

28 Ohno, K. et al. Congenital myasthenic syndrome caused by decreased agonist binding affinity due to a mutation in the acetylcholine receptor epsilon subunit. Neuron 17, 157–170 (1996). 10.1016/s0896-6273(00)80289-5.

29 Engel, A. G., Shen, X. M., Selcen, D. & Sine, S. M. Congenital myasthenic syndromes: pathogenesis, diagnosis, and treatment. Lancet Neurol 14, 420–434 (2015). 10.1016/s1474-4422(14)70201-7.

30 Wang, Y. et al. Germline genetic patterns underlying familial rheumatoid arthritis, systemic lupus erythematosus and primary Sjögren’s syndrome highlight T cell-initiated autoimmunity. Ann Rheum Dis 79, 268–275 (2020). 10.1136/annrheumdis-2019-215533.

31 Tan, E. M. et al. The 1982 revised criteria for the classification of systemic lupus erythematosus. Arthritis Rheum 25, 1271–1277 (1982). 10.1002/art.1780251101.

32 Hochberg, M. C. Updating the American College of Rheumatology revised criteria for the classification of systemic lupus erythematosus. Arthritis Rheum 40, 1725 (1997). 10.1002/art.1780400928.

33 Petri, M. et al. Derivation and validation of the Systemic Lupus International Collaborating Clinics classification criteria for systemic lupus erythematosus. Arthritis Rheum 64, 2677–2686 (2012). 10.1002/art.34473.

34 Aringer, M. et al. 2019 European League Against Rheumatism/American College of Rheumatology classification criteria for systemic lupus erythematosus. Ann Rheum Dis 78, 1151–1159 (2019). 10.1136/annrheumdis-2018-214819.

35 Miyazaki, T., Dierich, A., Benoist, C. & Mathis, D. Independent modes of natural killing distinguished in mice lacking Lag3. Science 272, 405–408 (1996). 10.1126/science.272.5260.405.

36 Shiokawa, M. et al. Laminin 511 is a target antigen in autoimmune pancreatitis. Sci Transl Med 10 (2018). 10.1126/scitranslmed.aaq0997.

