## Supplementary Information for "A Novel Autoantibody against LAG-3 in Systemic Lupus Erythematosus"

### SUPPLEMENTARY FIGURES

**Supplementary Figure 1. Anti-LAG-3 autoantibodies in patients with SLE mainly belong to the IgG1 subclass.**

(a) Serum levels of anti-LAG-3 autoantibodies of IgG subclasses (IgG1, IgG2, IgG3, IgG4) were evaluated using ELISA in patients with SLE (n=40) and healthy controls (n=32).

(b) Serum levels of IgA, IgM, and IgE isotypes of anti-LAG-3 autoantibodies were assessed in the same cohorts. The dashed line in each panel indicates the cutoff value, defined as the mean optical density (OD) of healthy controls plus 3 standard deviations (mean + 3SD).

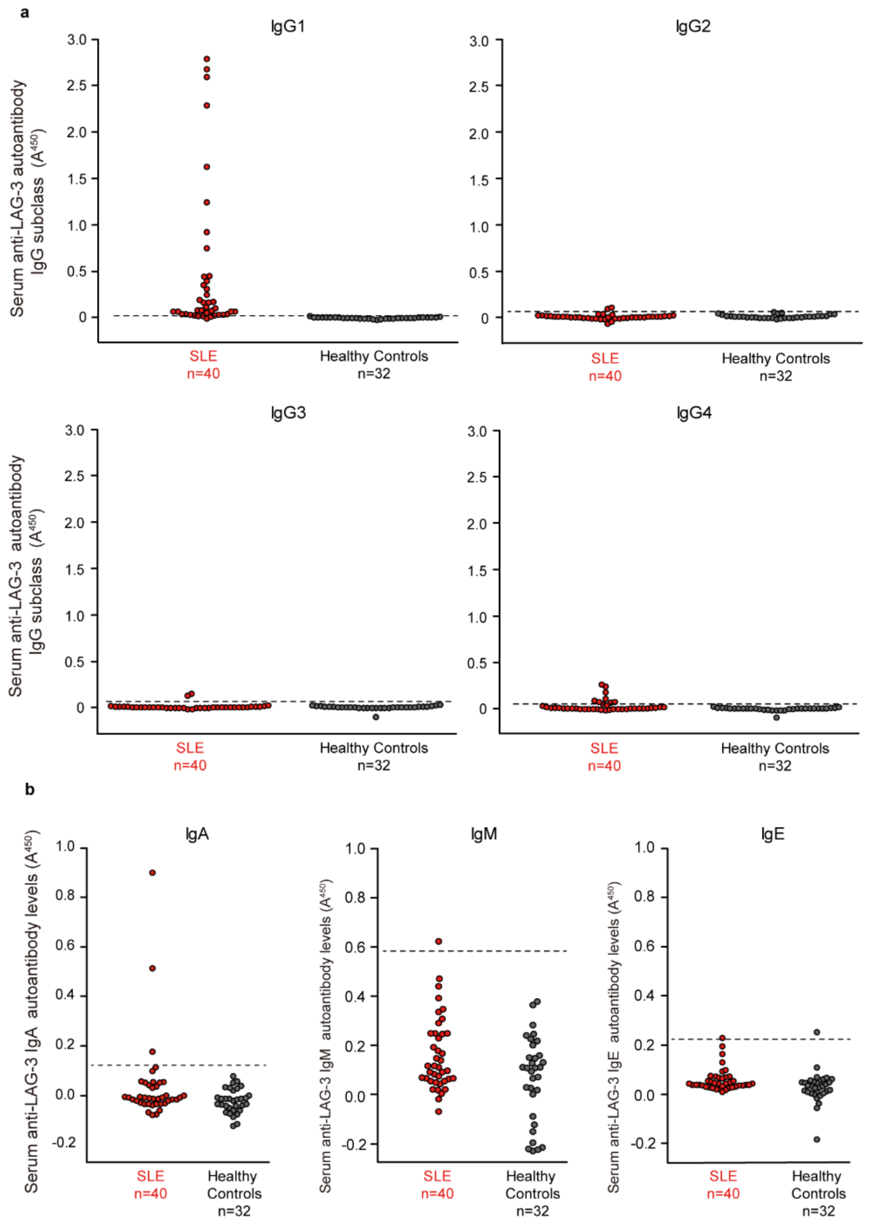

**Supplementary Figure 2. Autoimmune features in aged male *Lag*3 knockout mice.**

(a) Percentage of antinuclear antibody (ANA)-positive male mice at 6, 12, 18, and 24 months of age in wild-type (WT) and *Lag3* knockout (*Lag3* KO) groups. Numbers above the bars indicate the number of ANA-positive mice over the total number of mice assessed at each age.

(b) Serum anti-dsDNA IgG levels in male WT and *Lag3* KO mice at each age, measured using ELISA.

n.s. = not significant. Asterisks indicate statistical significance (*P<0.05) as determined using Fisher's exact test (panel a) or the Mann–Whitney U test (panel b). For box plots, centre lines indicate medians; boxes indicate the IQR; whiskers extend to 1.5× IQR; data points beyond whiskers are shown individually.

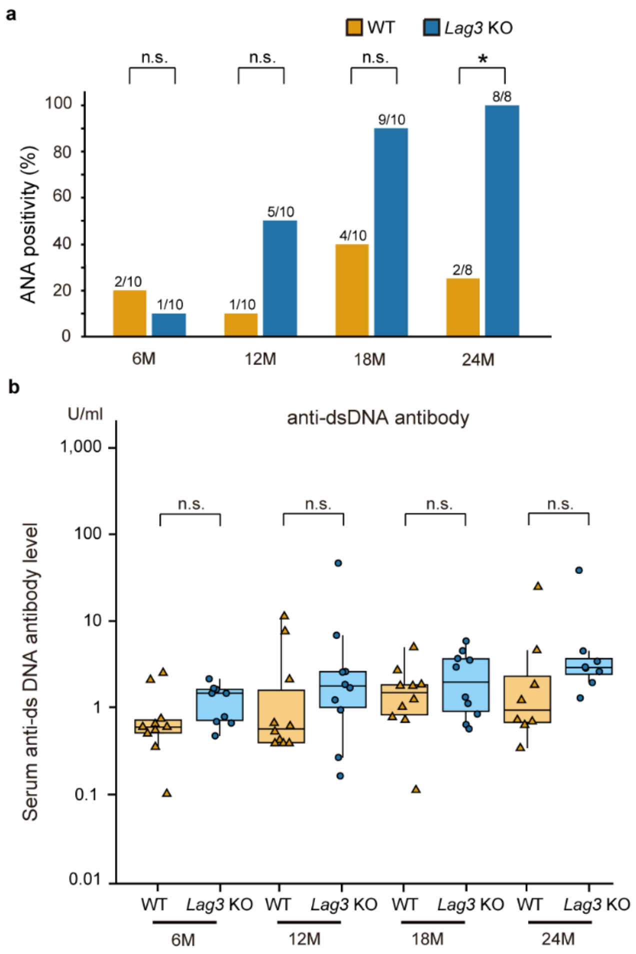

### SUPPLEMENTARY TABLES

#### Supplementary Table 1. Clinical characteristics of patients with systemic lupus erythematosus

| Patien t ID | Ag e | Sex | SLEDAI-2K | anti-dsDNA antibody positive | anti-Sm antibody positive | Low C3 | Low C4 | Low CH50 | IgG (mg/dl) | Lupus Nephritis | [a]Lupus nephritis class (ISN/RPS) | Other Clinical Features | anti-LAG-3 autoantibody positive |
|---|---|---|---|---|---|---|---|---|---|---|---|---|---|
| SLE 1 | 48 | F | 6 | Yes | No | No | No | No | 3695 | No | N/A | Leukopenia, Thrombocytopenia | No |
| SLE 2 | 48 | M | 1 | No | Yes | No | No | No | 687 | No | N/A | Fever, Arthritis, Neuropsychiatric SLE, Leukopenia, Thrombocytopenia | No |
| SLE 3 | 68 | F | 2 | No | Yes | Yes | No | No | 887 | No | N/A | Malar rash, Arthritis, Serositis, Neuropsychiatric SLE, Leukopenia, Thrombocytopenia | No |
| SLE 4 | 20 | F | 1 | No | No | Yes | No | No | 753 | Yes | V | Malar rash, Photosensitivity, Neuropsychiatric SLE, Leukopenia | Yes |
| SLE 5 | 28 | F | 2 | Yes | No | No | No | No | 1594 | No | N/A | Fever, Malar rash, Alopecia, Photosensitivity, Arthritis, Leukopenia, Thrombocytopenia | Yes |
| SLE 6 | 39 | F | 2 | No | Yes | Yes | No | No | 1847 | Yes | V | Fever, Mucosal ulcers, Malar rash, Photosensitivity, Arthritis | Yes |
| SLE 7 | 31 | F | 23 | Yes | Yes | Yes | Yes | Yes | 729 | Yes | IV-S (A) | Fever, Malar rash, Alopecia, Arthritis, Thrombocytopenia | No |
| SLE 8 | 22 | F | 8 | Yes | Yes | Yes | Yes | Yes | 2221 | Yes | V | Malar rash, Alopecia, Photosensitivity, Leukopenia | Yes |

| | | | | | | | | | | | | | |
|---|---|---|---|---|---|---|---|---|---|---|---|---|---|
| SLE 9 | 24 | F | 11 | Yes | Yes | Yes | No | No | 1664 | No | N/A | Fever, Mucosal ulcers, Malar rash, Photosensitivity, Arthritis, Neuropsychiatric SLE, Leukopenia | Yes |
| SLE 10 | 36 | M | 16 | Yes | Yes | Yes | Yes | Yes | 1208 | Yes | NE | Alopecia, Arthritis | Yes |
| SLE 11 | 39 | F | 23 | Yes | Yes | Yes | Yes | Yes | 3057 | No | N/A | Fever, Mucosal ulcers, Malar rash, Alopecia, Photosensitivity, Arthritis, Neuropsychiatric SLE, Leukopenia, Thrombocytopenia | No |
| SLE 12 | 25 | M | 14 | Yes | No | No | No | No | 3919 | Yes | NE | Fever, Serositis, Leukopenia, Thrombocytopenia | No |
| SLE 13 | 37 | F | 2 | Yes | Yes | No | No | No | 3002 | No | N/A | Fever, Malar rash, Alopecia, Arthritis, Leukopenia | No |
| SLE 14 | 32 | F | 17 | Yes | No | Yes | Yes | Yes | 1916 | Yes | III (A) + V | Leukopenia | Yes |
| SLE 15 | 45 | F | 6 | Yes | Yes | No | Yes | No | 3592 | No | N/A | Fever, Malar rash, Arthritis | No |
| SLE 16 | 38 | F | 4 | Yes | No | Yes | No | Yes | 1409 | Yes | III (A) | Fever, Alopecia, Arthritis | Yes |
| SLE 17 | 18 | F | 4 | No | Yes | No | No | No | 558 | No | N/A | Fever, Malar rash, Photosensitivity, Arthritis, Neuropsychiatric SLE, Leukopenia | No |
| SLE 18 | 56 | F | 0 | Yes | No | No | No | No | 1243 | Yes | NE | Malar rash, Photosensitivity, Arthritis | Yes |
| SLE 19 | 31 | F | 2 | Yes | No | No | No | No | 954 | Yes | NE | Fever, Malar rash, Alopecia, Photosensitivity, Arthritis, Serositis, Leukopenia, Thrombocytopenia | Yes |
| SLE 20 | 58 | F | 11 | Yes | No | Yes | No | No | 476 | Yes | NE | Serositis, Leukopenia, Thrombocytopenia | Yes |

| | | | | | | | | | | | | | |
|---|---|---|---|---|---|---|---|---|---|---|---|---|---|
| SLE 21 | 41 | F | 4 | Yes | No | Yes | Yes | Yes | 3456 | No | N/A | Fever, Alopecia, Photosensitivity, Arthritis, Serositis, Neuropsychiatric SLE, Leukopenia, Thrombocytopenia | No |
| SLE 22 | 39 | F | 2 | No | No | Yes | No | Yes | 960 | No | N/A | Fever, Mucosal ulcers, Malar rash, Alopecia, Photosensitivity, Arthritis | No |
| SLE 23 | 80 | F | 2 | Yes | No | No | No | No | 1801 | No | N/A | Malar rash, Photosensitivity | No |
| SLE 24 | 26 | M | 12 | Yes | Yes | Yes | Yes | Yes | 2175 | Yes | II | Fever, Malar rash, Alopecia, Serositis, Leukopenia, Thrombocytopenia | No |
| SLE 25 | 37 | F | 4 | No | No | No | No | No | 544 | Yes | IV-G (A/C) + V | Malar rash, Alopecia | Yes |
| SLE 26 | 30 | M | 0 | No | No | No | No | No | 1476 | No | N/A | Fever, Malar rash, Alopecia, Photosensitivity, Arthritis, Leukopenia | No |
| SLE 27 | 62 | M | 0 | No | No | No | No | No | 830 | No | N/A | Fever, Mucosal ulcers, Malar rash, Alopecia, Photosensitivity, Arthritis, Leukopenia, Thrombocytopenia | No |
| SLE 28 | 72 | F | 0 | No | No | Yes | No | No | 1865 | No | N/A | Malar rash, Arthritis, Leukopenia | Yes |
| SLE 29 | 44 | F | 0 | No | Yes | No | No | No | 1075 | Yes | III (C) | Fever, Malar rash, Alopecia, Neuropsychiatric SLE | No |
| SLE 30 | 52 | M | 1 | Yes | No | Yes | Yes | Yes | 1030 | Yes | NE | Arthritis, Neuropsychiatric SLE, Leukopenia, Thrombocytopenia | No |
| SLE 31 | 31 | F | 1 | Yes | Yes | Yes | No | Yes | 921 | Yes | III (A) + V | Fever, Malar rash, Alopecia, Photosensitivity, | No |

| | | | | | | | | | | | | | |
|---|---|---|---|---|---|---|---|---|---|---|---|---|---|
| | | | | | | | | | | | | Arthritis, Serositis, Leukopenia | |
| SLE 32 | 50 | F | 0 | No | No | No | No | No | 821 | No | N/A | Fever, Malar rash, Alopecia, Photosensitivity, Arthritis, Serositis, Leukopenia, Thrombocytopenia | No |
| SLE 33 | 77 | M | 1 | No | No | No | No | No | 1034 | No | N/A | Fever, Photosensitivity, Arthritis, Leukopenia, Thrombocytopenia | No |
| SLE 34 | 45 | F | 0 | No | No | No | No | No | 1416 | No | N/A | Mucosal ulcers, Alopecia, Photosensitivity, Arthritis | Yes |
| SLE 35 | 25 | F | 0 | No | No | No | No | No | 863 | Yes | NE | Alopecia, Leukopenia | No |
| SLE 36 | 75 | F | 0 | Yes | No | No | No | No | 1066 | Yes | III (A) | Arthritis | No |
| SLE 37 | 37 | F | 8 | Yes | Yes | Yes | Yes | Yes | 1370 | Yes | III (A/C) + V | Malar rash, Photosensitivity, Arthritis | No |
| SLE 38 | 36 | F | 6 | Yes | No | No | No | No | 1760 | Yes | NE | Fever, Alopecia, Arthritis | Yes |
| SLE 39 | 48 | F | 0 | No | No | No | No | No | 1265 | Yes | NE | Fever, Photosensitivity, Arthritis, Leukopenia | No |
| SLE 40 | 35 | F | 2 | No | No | Yes | No | No | 656 | Yes | II | Fever, Mucosal ulcers, Malar rash, Alopecia, Photosensitivity, Arthritis, Leukopenia | No |
| SLE 41 | 45 | F | 2 | Yes | No | Yes | No | No | 1013 | Yes | IV (A) | Fever, Alopecia, Arthritis | Yes |
| SLE 42 | 28 | M | 4 | Yes | No | Yes | Yes | Yes | 849 | Yes | IV-G (A/C) | Fever, Alopecia, Neuropsychiatric SLE | Yes |
| SLE 43 | 38 | F | 0 | No | No | No | No | No | 1609 | No | N/A | Fever, Malar rash, Alopecia, Photosensitivity, Arthritis, Leukopenia, Thrombocytopenia | No |
| SLE 44 | 35 | F | 5 | Yes | No | Yes | No | No | 1074 | No | N/A | Arthritis, Serositis, Leukopenia, Thrombocytopenia | No |
| SLE 45 | 40 | F | 0 | No | No | No | No | No | 995 | Yes | NE | Fever, Photosensitivity, Arthritis | Yes |

| | | | | | | | | | | | | | |
|---|---|---|---|---|---|---|---|---|---|---|---|---|---|
| SLE 46 | 58 | F | 3 | Yes | No | No | No | No | 1478 | No | N/A | Fever, Malar rash, Arthritis, Serositis, Leukopenia, Thrombocytopenia | No |
| SLE 47 | 44 | F | 0 | No | No | No | No | No | 639 | No | N/A | Neuropsychiatric SLE | No |
| SLE 48 | 68 | F | 0 | No | No | No | No | No | 554 | No | N/A | Fever, Malar rash, Photosensitivity, Neuropsychiatric SLE, Leukopenia, Thrombocytopenia | No |
| SLE 49 | 30 | F | 2 | Yes | No | No | No | No | 1172 | No | N/A | Fever, Malar rash, Alopecia, Photosensitivity, Arthritis, Neuropsychiatric SLE, Leukopenia, Thrombocytopenia | No |
| SLE 50 | 42 | F | 4 | No | Yes | No | No | No | 1419 | Yes | IV | Malar rash, Leukopenia | Yes |
| SLE 51 | 41 | F | 2 | Yes | No | No | No | No | 786 | Yes | IV-S (A/C) | Fever, Malar rash, Alopecia, Arthritis, Serositis, Leukopenia | No |
| SLE 52 | 59 | F | 2 | Yes | No | Yes | No | No | 3502 | No | N/A | Arthritis | No |
| SLE 53 | 53 | F | 6 | Yes | No | No | No | No | 1282 | No | N/A | Mucosal ulcers, Malar rash, Alopecia, Photosensitivity, Arthritis, Leukopenia | No |
| SLE 54 | 56 | F | 0 | Yes | Yes | No | No | No | 692 | Yes | IV-G (A/C) | Fever, Arthritis, Serositis | Yes |
| SLE 55 | 59 | F | 0 | Yes | No | No | No | No | 1440 | Yes | IV-G (A) | Fever, Leukopenia | Yes |
| SLE 56 | 41 | F | 2 | Yes | Yes | No | No | No | 1062 | No | N/A | Fever, Malar rash, Neuropsychiatric SLE, Leukopenia | No |
| SLE 57 | 32 | F | 0 | No | No | Yes | No | No | 1362 | Yes | II | Fever, Malar rash, Alopecia, Photosensitivity, Arthritis, Leukopenia | No |
| SLE 58 | 48 | F | 3 | No | Yes | Yes | Yes | No | 2231 | No | N/A | Fever, Malar rash, Alopecia, Photosensitivity, | No |

| | | | | | | | | | | | | | |
|---|---|---|---|---|---|---|---|---|---|---|---|---|---|
| | | | | | | | | | | | | Arthritis, Leukopenia, Thrombocytopenia | |
| SLE 59 | 71 | F | 0 | No | No | No | No | No | 902 | No | N/A | Mucosal ulcers, Malar rash, Alopecia, Photosensitivity, Arthritis, Leukopenia, Thrombocytopenia | No |
| SLE 60 | 50 | F | 4 | Yes | No | No | No | No | 972 | Yes | III (A/C) | Fever, Arthritis, Leukopenia | Yes |
| SLE 61 | 26 | F | 6 | Yes | No | No | No | No | 872 | Yes | III (A/C) + V | Fever, Malar rash, Photosensitivity, Arthritis, Leukopenia | Yes |
| SLE 62 | 36 | F | 2 | Yes | No | No | No | No | 1392 | No | N/A | Fever, Malar rash, Photosensitivity, Arthritis, Serositis, Leukopenia, Thrombocytopenia | No |
| SLE 63 | 71 | F | 2 | Yes | No | No | No | No | 2108 | No | N/A | Malar rash, Arthritis, Neuropsychiatric SLE, Leukopenia, Thrombocytopenia | No |
| SLE 64 | 30 | F | 2 | Yes | No | Yes | No | No | 990 | No | N/A | Fever, Mucosal ulcers, Malar rash, Alopecia, Photosensitivity, Leukopenia | No |
| SLE 65 | 54 | F | 0 | No | No | No | No | No | 949 | Yes | IV-G (A) + V | Mucosal ulcers, Arthritis, Leukopenia, Thrombocytopenia | No |
| SLE 66 | 39 | F | 2 | No | No | Yes | No | No | 1044 | Yes | NE | Alopecia, Photosensitivity, Leukopenia, Thrombocytopenia | No |
| SLE 67 | 55 | F | 0 | Yes | No | No | No | No | 1056 | No | N/A | Fever | No |
| SLE 68 | 56 | F | 0 | Yes | No | No | No | No | 1374 | No | N/A | Fever, Mucosal ulcers, Malar rash, Arthritis, Neuropsychiatric SLE | No |
| SLE 69 | 53 | F | 4 | Yes | No | Yes | No | No | 1230 | No | N/A | Fever, Arthritis, Leukopenia | No |

| | | | | | | | | | | | | | |
|---|---|---|---|---|---|---|---|---|---|---|---|---|---|
| SLE 70 | 60 | F | 0 | No | No | No | No | No | 1401 | No | N/A | Malar rash, Photosensitivity, Arthritis | No |
| SLE 71 | 31 | M | 5 | Yes | No | No | Yes | Yes | 987 | Yes | IV-G (A/C) | Fever, Mucosal ulcers, Malar rash, Alopecia, Photosensitivity, Arthritis, Leukopenia | Yes |
| SLE 72 | 36 | F | 0 | Yes | No | No | No | No | 1116 | No | N/A | Fever, Malar rash, Alopecia, Photosensitivity, Arthritis, Leukopenia, Thrombocytopenia | No |
| SLE 73 | 56 | F | 0 | Yes | No | No | No | No | 1446 | No | N/A | Fever, Malar rash, Photosensitivity, Arthritis, Neuropsychiatric SLE | No |
| SLE 74 | 30 | F | 2 | Yes | Yes | Yes | No | No | 864 | No | N/A | Fever, Malar rash, Alopecia, Photosensitivity, Arthritis | No |
| SLE 75 | 52 | F | 0 | No | Yes | No | No | No | 1614 | No | N/A | Fever, Alopecia, Arthritis, Leukopenia, Thrombocytopenia | No |
| SLE 76 | 53 | F | 2 | Yes | No | No | No | No | 1217 | No | N/A | Arthritis, Leukopenia | Yes |
| SLE 77 | 46 | F | 6 | Yes | No | No | No | No | 949 | Yes | NE | Fever, Malar rash | Yes |
| SLE 78 | 47 | F | 0 | No | No | No | No | No | 1067 | Yes | III | Fever, Photosensitivity, Arthritis, Neuropsychiatric SLE, Leukopenia | Yes |
| SLE 79 | 52 | F | 3 | No | Yes | No | No | No | 1824 | No | N/A | Fever, Malar rash, Arthritis, Leukopenia | Yes |
| SLE 80 | 44 | F | 3 | Yes | Yes | No | No | No | 1992 | Yes | V | Arthritis, Thrombocytopenia | No |
| SLE 81 | 72 | F | 0 | No | Yes | No | No | No | 1021 | No | N/A | Mucosal ulcers, Malar rash, Photosensitivity, Arthritis, Leukopenia | No |
| SLE 82 | 49 | F | 4 | No | NE | Yes | Yes | Yes | 1131 | No | N/A | Malar rash, Arthritis, Serositis | No |
| SLE 83 | 79 | F | 0 | No | No | No | No | No | 1502 | No | N/A | Mucosal ulcers, Malar rash, Photosensitivity, Arthritis, Leukopenia | No |

| | | | | | | | | | | | | | |
|---|---|---|---|---|---|---|---|---|---|---|---|---|---|
| SLE 84 | 59 | F | 0 | No | NE | No | No | No | 1132 | Yes | NE | Photosensitivity | Yes |
| SLE 85 | 73 | F | 0 | No | NE | No | No | No | 1653 | No | N/A | Mucosal ulcers | No |
| SLE 86 | 81 | M | 0 | Yes | No | No | No | No | 1370 | Yes | NE | Malar rash, Arthritis, Neuropsychiatric SLE | No |
| SLE 87 | 47 | F | 0 | No | Yes | No | No | No | 1121 | No | N/A | Fever, Mucosal ulcers, Malar rash, Alopecia, Photosensitivity, Arthritis, Serositis, Neuropsychiatric SLE, Leukopenia, Thrombocytopenia | No |
| SLE 88 | 33 | F | 4 | Yes | Yes | No | Yes | Yes | 1728 | No | N/A | Fever, Malar rash, Alopecia, Photosensitivity, Arthritis, Leukopenia, Thrombocytopenia | No |
| SLE 89 | 43 | F | 5 | No | No | No | No | No | 1396 | No | N/A | Arthritis, Serositis | Yes |
| SLE 90 | 77 | F | 0 | No | No | No | No | No | 1305 | Yes | NE | Fever, Alopecia, Photosensitivity, Arthritis, Serositis, Leukopenia | No |
| SLE 91 | 59 | F | 0 | No | Yes | No | No | No | 991 | No | N/A | Fever, Malar rash, Alopecia, Photosensitivity, Arthritis | Yes |
| SLE 92 | 48 | F | 0 | No | Yes | No | No | No | 1150 | No | N/A | Malar rash, Photosensitivity, Arthritis, Leukopenia | Yes |
| SLE 93 | 25 | F | 2 | No | Yes | No | Yes | No | 1406 | No | N/A | Fever, Arthritis, Serositis | Yes |
| SLE 94 | 34 | F | 2 | No | No | Yes | No | Yes | 1298 | No | N/A | Fever, Malar rash, Photosensitivity, Arthritis, Leukopenia | No |
| SLE 95 | 49 | M | 18 | Yes | Yes | Yes | Yes | Yes | 2837 | Yes | V | Malar rash, Alopecia, Arthritis, Leukopenia | No |
| SLE 96 | 31 | F | 2 | Yes | No | No | No | No | 1860 | No | N/A | Fever, Mucosal ulcers, Malar rash, Arthritis, Serositis, Leukopenia | Yes |
| SLE 97 | 66 | F | 0 | No | Yes | No | No | No | 1720 | Yes | NE | Malar rash, Neuropsychiatric SLE | Yes |

| | | | | | | | | | | | | | |
|---|---|---|---|---|---|---|---|---|---|---|---|---|---|
| SLE 98 | 39 | F | 0 | No | No | No | No | No | 1177 | Yes | II | Arthritis | Yes |
| SLE 99 | 65 | F | 0 | No | No | No | No | No | 999 | No | N/A | Serositis | No |
| SLE 100 | 47 | F | 0 | Yes | Yes | No | No | No | 1894 | No | N/A | Alopecia, Photosensitivity, Arthritis, Serositis, Leukopenia | Yes |
| SLE 101 | 52 | F | 5 | Yes | Yes | No | Yes | No | 1036 | No | N/A | Fever, Alopecia, Photosensitivity, Arthritis, Leukopenia, Thrombocytopenia | Yes |
| SLE 102 | 28 | F | 11 | Yes | No | Yes | Yes | Yes | 1330 | No | N/A | Malar rash, Leukopenia | Yes |
| SLE 103 | 21 | F | 23 | Yes | No | Yes | Yes | Yes | 1557 | No | N/A | Fever, Malar rash, Arthritis, Leukopenia | Yes |
| SLE 104 | 37 | F | 6 | No | Yes | No | No | Yes | 2001 | No | N/A | Fever, Malar rash, Alopecia, Arthritis, Leukopenia, Thrombocytopenia | Yes |
| SLE 105 | 29 | F | 2 | No | No | No | No | No | 919 | Yes | III (A) | Malar rash, Alopecia, Photosensitivity, Arthritis, Leukopenia, Thrombocytopenia | No |
| SLE 106 | 22 | M | 4 | Yes | Yes | No | No | Yes | 1891 | Yes | III (A) | Fever, Mucosal ulcers, Leukopenia, Thrombocytopenia | No |
| SLE 107 | 71 | F | 8 | Yes | No | Yes | Yes | No | 2278 | Yes | IV-G (A/C) | Fever, Arthritis, Leukopenia, Thrombocytopenia | No |
| SLE 108 | 50 | F | 14 | Yes | Yes | Yes | Yes | Yes | 1748 | Yes | III (A/C) | Fever, Malar rash, Alopecia, Photosensitivity, Arthritis, Serositis, Leukopenia, Thrombocytopenia | Yes |
| SLE 109 | 55 | F | 0 | Yes | Yes | No | No | No | 1420 | No | N/A | Fever, Arthritis, Serositis | Yes |
| SLE 110 | 54 | F | 3 | Yes | No | No | No | No | 2503 | No | N/A | Alopecia, Arthritis, Serositis, Leukopenia | Yes |

| | | | | | | | | | | | | | |
|---|---|---|---|---|---|---|---|---|---|---|---|---|---|
| SLE 111 | 76 | F | 0 | No | No | No | No | No | 835 | Yes | IV | Fever, Mucosal ulcers, Alopecia, Photosensitivity, Arthritis | No |
| SLE 112 | 53 | F | 0 | No | No | No | No | No | 1970 | No | N/A | Fever, Alopecia, Photosensitivity, Leukopenia | No |
| SLE 113 | 46 | F | 14 | Yes | Yes | Yes | Yes | Yes | 1466 | Yes | NE | Fever, Serositis, Leukopenia | No |
| SLE 114 | 23 | F | 17 | Yes | Yes | Yes | Yes | Yes | 3065 | Yes | IV-G (A) | Fever, Mucosal ulcers, Leukopenia, Thrombocytopenia | Yes |
| SLE 115 | 32 | F | 2 | Yes | Yes | Yes | No | Yes | 1058 | No | N/A | Fever, Mucosal ulcers, Malar rash, Alopecia, Photosensitivity, Arthritis, Leukopenia | No |
| SLE 116 | 77 | F | 2 | Yes | Yes | No | No | No | 1363 | Yes | II | Fever, Photosensitivity, Leukopenia | No |
| SLE 117 | 57 | F | 0 | Yes | Yes | No | No | No | 976 | Yes | V | Fever, Alopecia, Arthritis, Leukopenia | Yes |
| SLE 118 | 21 | F | 14 | Yes | Yes | Yes | Yes | Yes | 2132 | Yes | V | Fever, Mucosal ulcers, Malar rash, Photosensitivity, Arthritis, Leukopenia, Thrombocytopenia | No |
| SLE 119 | 67 | F | 1 | No | No | No | No | No | 728 | Yes | V | Fever, Mucosal ulcers, Arthritis, Neuropsychiatric SLE, Leukopenia | Yes |
| SLE 120 | 43 | F | 17 | Yes | No | Yes | Yes | Yes | 1742 | Yes | IV-G (A) | Leukopenia, Thrombocytopenia | No |
| SLE 121 | 53 | F | 8 | Yes | No | Yes | Yes | Yes | 4256 | No | N/A | Fever, Mucosal ulcers, Arthritis, Thrombocytopenia | No |
| SLE 122 | 54 | F | 18 | Yes | Yes | Yes | Yes | Yes | 2809 | Yes | NE | Malar rash | Yes |
| SLE 123 | 39 | F | 2 | No | No | Yes | Yes | Yes | 1660 | No | N/A | Fever, Malar rash, Arthritis, Leukopenia | Yes |

| | | | | | | | | | | | | | |
|---|---|---|---|---|---|---|---|---|---|---|---|---|---|
| SLE 124 | 68 | F | 4 | Yes | No | No | No | No | 4554 | No | N/A | Arthritis, Neuropsychiatric SLE | No |
| SLE 125 | 21 | M | 20 | Yes | No | Yes | Yes | Yes | 435 | Yes | IV-S (A/C) + V | Malar rash, Neuropsychiatric SLE, Leukopenia | Yes |
| SLE 126 | 66 | F | 6 | Yes | No | No | No | No | 1242 | Yes | II | Fever, Alopecia, Photosensitivity, Arthritis, Neuropsychiatric SLE, Leukopenia | No |
| SLE 127 | 20 | F | 8 | Yes | Yes | No | Yes | Yes | 2109 | No | N/A | Malar rash, Photosensitivity, Arthritis | Yes |
| SLE 128 | 70 | M | 15 | Yes | No | Yes | No | Yes | 2656 | Yes | IV-S (A/C) | Fever, Arthritis, Serositis, Leukopenia, Thrombocytopenia | No |
| SLE 129 | 53 | F | 0 | No | No | No | No | No | 1049 | Yes | IV-G (A) | Fever, Malar rash, Photosensitivity, Thrombocytopenia | No |
| SLE 130 | 55 | F | 5 | No | No | No | No | No | 1851 | No | N/A | Malar rash, Alopecia, Photosensitivity, Arthritis, Leukopenia, Thrombocytopenia | Yes |

[a]Lupus nephritis was classified according to the ISN/RPS classification, where histological evaluation was available.

N/A, not applicable; NE, not evaluable

**Supplementary Table 2. Demographic and clinical characteristics of patients with autoimmune diseases other than SLE and anti-LAG-3 antibody positivity rates by disease.**

| Characteristic | n (%) or median (range) | Anti-LAG-3 autoantibody positive, n (%) |
|---|---|---|
| **Total** | 119 | 4 (3.4) |
| Age, median (range), years | 63 (15–95) | — |
| Female sex | 87 (73.1) | — |
| **Disease** | | |
| Dermatomyositis/Polymyositis | 39 (32.8) | 0 (0) |
| Systemic sclerosis | 22 (18.5) | 2 (9.1) |
| Rheumatoid arthritis | 22 (18.5) | 1 (4.5) |
| Sjögren's syndrome | 15 (12.6) | 1 (6.7) |
| Mixed connective tissue disease | 7 (5.9) | 0 (0) |
| Behçet's disease | 6 (5.0) | 0 (0) |
| Others | 8 (6.7) | 0 (0) |

**Supplementary Table 3. Clinical characteristics of serum samples used in the LAG-3/MHC class II blockade assay**

| Patient ID | Age | Sex | SLEDAI-2K | anti-ds DNA antibody positive | anti-Sm antibody positive | Low C3 | Low C4 | Low CH50 | Lupus Nephritis | Other Clinical Features | Time from initial diagnosis of SLE (mo) |
|---|---|---|---|---|---|---|---|---|---|---|---|
| SLE-pos A | 12 | F | 31 | Yes | No | Yes | Yes | Yes | Yes | Hemolytic anemia, Vasculitis, Rash, Alopecia, Mucosal ulcer, Photosensitivity, Fever | 0 |
| SLE-pos B | 8 | F | 18 | Yes | Yes | Yes | Yes | Yes | Yes | Leukopenia, Hemolytic anemia, Vasculitis, Rash, Mucosal ulcer, Fever | 2 |
| SLE-pos C | 15 | M | 23 | Yes | No | Yes | Yes | Yes | No | Hemolytic anemia, Myositis, Arthritis, Rash, Fever | 1 |
| SLE-pos D | 9 | M | 12 | Yes | Yes | Yes | Yes | Yes | Yes | Rash, Pericarditis | 1 |
| SLE-pos E | 10 | F | 13 | Yes | No | Yes | Yes | Yes | Yes | Arthritis, Fever | 0 |
| SLE-pos F | 33 | F | 9 | Yes | Yes | No | Yes | No | No | Leukopenia, Anemia, Pericarditis | 1 |
| SLE-neg A | 46 | F | 14 | Yes | Yes | Yes | Yes | Yes | Yes | Fever, Serositis, Leukopenia | 2 |
| SLE-neg B | 70 | M | 15 | Yes | No | Yes | No | Yes | Yes | Fever, Arthritis, Serositis, Leukopenia, Thrombocytopenia | 0 |
| SLE-neg C | 43 | F | 17 | Yes | No | Yes | Yes | No | Yes | Leukopenia, Thrombocytopenia | 2 |
| SLE-neg D | 11 | F | 18 | Yes | No | Yes | Yes | Yes | Yes | Alopecia | 4 |
| SLE-neg E | 13 | F | 26 | Yes | Yes | Yes | Yes | Yes | Yes | Vasculitis, Arthritis, Rash, Alopecia, Mucosal ulcer, Photosensitivity, Serositis, Pericarditis | 6 |
| SLE-neg F | 14 | F | 23 | Yes | No | No | No | No | Yes | Vasculitis, Arthritis, Rash, Hemolytic anemia, Fever | 0 |
